# PCAGP, a parallel convolutional attention network-based method for crop genomic prediction

**DOI:** 10.1101/2025.05.17.654636

**Authors:** Wenyu Peng, Yuxia Sheng, Ying Zhou, Li Chai

## Abstract

Genomic selection (GS) has emerged as a transformative breeding paradigm, leveraging genome-wide marker data to predict phenotypic outcomes and accelerate crop improvement cycles. Traditional statistical models face fundamental limitations in modeling the higher-order, non-linear genotype-phenotype relationships inherent in complex agronomic traits. Deep learning architectures have recently shown remarkable predictive performance in GS applications. Current deep learning approaches in GS predominantly employ a serial structure convolution kernel, this may lead to gradient vanishing and computational efficiency challenges. In this study, we propose a method based on a parallel convolutional attention network for crop genomic prediction (PCAGP), a novel deep learning framework for crop genomic prediction. The architecture uses a convolution kernel to transmit information through a parallel structure, combined with a coordinate attention mechanism that effectively integrates inter-channel relationships and spatial genomic information. This approach significantly improves the accuracy of phenotypic prediction. We compared PCAGP with six widely used GS methods on four benchmark datasets, including statistical method (GBLUP), machine learning approaches (SVR, RF), and deep learning models (DeepGS, DNNGP, and SoyDNGP). The experimental results demonstrate that PCAGP outperforms all baseline methods, and the average prediction accuracy is improved (0.4%-53.1%).

## Introduction

As uncertainty about climate change intensifies, the issue of food security becomes more acute. Adverse factors such as extreme weather, the frequency of pests and diseases, and the increasing scarcity of land resources will further threaten crop yields and stability (He et al., 2020). In order to reduce overuse of land resources and cope with the sharp increase in food demand in the future, the development of high-yield and disease resistant crop varieties has become a key task (Ewert et al., 2023). Through accurate prediction of crop traits, breeders can select varieties with superior agronomic traits more effectively. This accelerates the promotion of new varieties to address the challenges caused by food shortages (Bali and Singla, 2022).

Genomic selection (GS) is an advanced breeding technology that has been widely applied in animal and plant breeding and demonstrates significant potential (Meuwissen et al., 2001). It is a breeding method that uses information from whole genome markers to predict the genetic potential of individuals. Compared to traditional breeding methods, GS does not require the identification of specific loci significantly associated with target traits. Instead, it captures the subtle effects of each locus in the genome through high-density genetic markers (Jannink et al., 2010). This ability makes GS have obvious advantages in capturing the genetic effects of complex traits, effectively improving crop yield and disease resistance. However, the effectiveness of GS is influenced by various factors, including the size of the training population, the heritability of traits, marker density, and the predictive methods employed (Wang et al., 2024).

Traditional statistical methods, such as genomic best linear unbiased prediction (GBLUP) (VanRaden, 2008) and Bayesian approaches (Nazzicari and Biscarini, 2022), have been widely used to model genotype effects and phenotype prediction. However, these models typically assume that the random effects of genotypes follow a certain prior distribution. In practical applications, the effects of genotypes are often unknown, and it is challenging to capture complex non-additive effects (Govaichelvan et al., 2024). For example, the GBLUP model assumes that all markers contribute equally to the relationship matrix, which may not be the case in reality. To address these limitations, several improved GBLUP-based methods have been developed. The GA-GBLUP method selects markers related to the target trait by introducing a genetic algorithm (GA) to optimize the weight (Xu et al., 2024). The multi-trait GBLUP model, which combines global or local genetic relevant information, is also used to improve the accuracy of multi-population genome prediction (Teng et al., 2024). Furthermore, the application of the L2,1 norm regularized multiple regression model in genomic selection for multiple traits provides new ideas for more efficient prediction of multiple traits (Mbebi et al., 2021). Beyond classical statistical methods, machine learning approaches such as random forest (RF) (Adetunji et al., 2022) and support vector regression (SVR) (Maenhout et al., 2007) have been introduced into the field of predicting crop agronomic traits (Prasad et al., 2021). Although these methods excel at capturing non-linear relationships, their performance can be constrained when handling high-dimensional and complex genomic datasets, highlighting the need for more advanced computational frameworks.

In recent years, deep learning (DL) methods have gained popularity in GS, facilitating the application of large-scale genomic data in phenotypic prediction (Xie et al., 2016). Researchers are able to identify genetic markers associated with traits more precisely, thereby accelerating traditional breeding processes and reducing costs (Beyene et al., 2021). Among these, convolution neural networks (CNNs) have shown significant advantages in the field of genomic selection, particularly in capturing complex nonlinear relationships and gene interactions. Genomic prediction model based on deep learning framework (DeepGS) (Ma et al., 2018), the deep neural network-based method for genomic prediction using multi-omics data in plants (DNNGP) (Wang et al., 2023), a network deep learning framework genomic prediction in soybean breeding (SoyDNGP) (Gao et al., 2023), and multi-modal deep learning models that integrate genome-wide genetic variation with rich phenotype observations to predict crop yield (PheGeMIL) (Togninalli et al., 2023), have made significant progress in crop breeding. However, these models all rely on serial-structured convolution kernels to extract features, which can lead to information loss when analyzing complex relationships within genomic data. Additionally, determining the optimal size of the convolution kernel remains a time-consuming and computationally intensive task. In contrast, a parallel convolution structure extracts features simultaneously through multiple convolution kernels of different sizes without requiring hyperparameter tuning, thereby improving feature representation capability and diversity. In addition, this structure has been shown to improve computational efficiency while preserving more details (Xie et al., 2024).

In this study, we propose a novel crop genomic prediction method based on a parallel convolutional attention network (PCAGP). The PCAGP model consists of a parallel convolution layer, a coordinate attention (CA) layer, and two fully connected (FC) layers. The parallel structure convolution layer extracts genotype features using convolution kernels of different sizes simultaneously. The CA layer effectively captures long-range dependencies while preserving precise positional information. The FC layer performs global feature fusion and extraction through linear transformations and nonlinear activation functions. To evaluate the performance of the PCAGP method, we conducted comprehensive benchmark experiments on datasets of soybean, wheat, and rice. The results demonstrate that, compared to six widely used prediction models, PCAGP achieves superior prediction accuracy for agronomic traits across different crop species.

## Materials and Methods

We used four real datasets to evaluate the performance of the proposed model.

### Soybean datasets

In this study, two soybean datasets were used. Soybean13784, originates from two integrated online databases, SoyBase and GRIN-Global (Grant et al., 2010; Postman et al., 2009). It contains genotypic data for 20087 soybean accessions based on the SoySNP50K iSelect BeadChip platform, including 42509 high-confidence single nucleotide polymorphisms (SNPs). After a selection process, 13784 samples and 32032 SNPs were used for the experiments. The corresponding selection of four phenotypic data is from the GRIN-Global database (https://npgsweb.arsgrin.gov/gringlobal/search): protein content (protein), oil content (oil), 100 seed weight (SdWgt), and yield (Yield).

The second dataset, soybean573, originates from the Yangtze-Huaihe Soybean Breeding Germplasm Population (YHSBLP) established by the National Soybean Improvement Center. It includes SNP data obtained through the simplified genome sequencing technology of RAD-seq (Karikari et al., 2020). This dataset includes genotyping information for 573 samples with a total of 61166 SNPs. In addition, it includes phenotypic data for the average weight of 100 seeds from the YHSBLP population, collected in 2013, 2014, 2017, and 2018 at the Jiangpu Experimental Station of Nanjing Agricultural University. The traits recorded are Yield 13: average hundred-seed weight in 2013, Yield 14: average hundred-seed weight in 2014, Yield 17: average hundred-seed weight in 2017, and Yield 18: average hundred-seed weight in 2018.

### Wheat dataset

The third dataset, wheat2000, consists of 2000 Iranian bread wheat landraces from the CIMMYT wheat genebank (Crossa et al., 2016). This dataset includes genotyping data for 33709 DArT markers from 2000 landraces and evaluations of six agronomic traits: grain length (GL), grain width (GW), grain hardness (GH), thousand kernel weight (TKW), test weight (TW), and grain protein (GP).

### Rice dataset

The fourth dataset, rice529, originates from Huazhong Agricultural University (Guo et al., 2018). It includes 17397026 SNPs from 529 rice samples. We used Plink software to remove SNPs with a deletion rate greater than 5% or a minor allele frequency (MAF) lower than 0.001, and then excluded the Hardy-Weinberg equilibrium test with a p value *<*1.0E-4, resulting in 742622 SNPs for subsequent analyzes. For this study, we used the 50000 SNPs that were most associated after screening. Four agronomic traits were included: stem dry weight (SDW), stem fresh weight (SFW), grain length (GL), and the ratio of total projected area to bounding rectangle area (TBR R).

### PCAGP model structure

Our proposed PCAGP is a deep learning model that consists of a parallel convolution layer, a coordinate attention mechanism, and fully connected layers. The parallel structure convolution layer is used to increase the width of the model and minimize the loss of information. The coordinate attention mechanism is a fusion of location and channel information. The architecture of PCAGP is shown in Figure 1.

**Figure 1.**
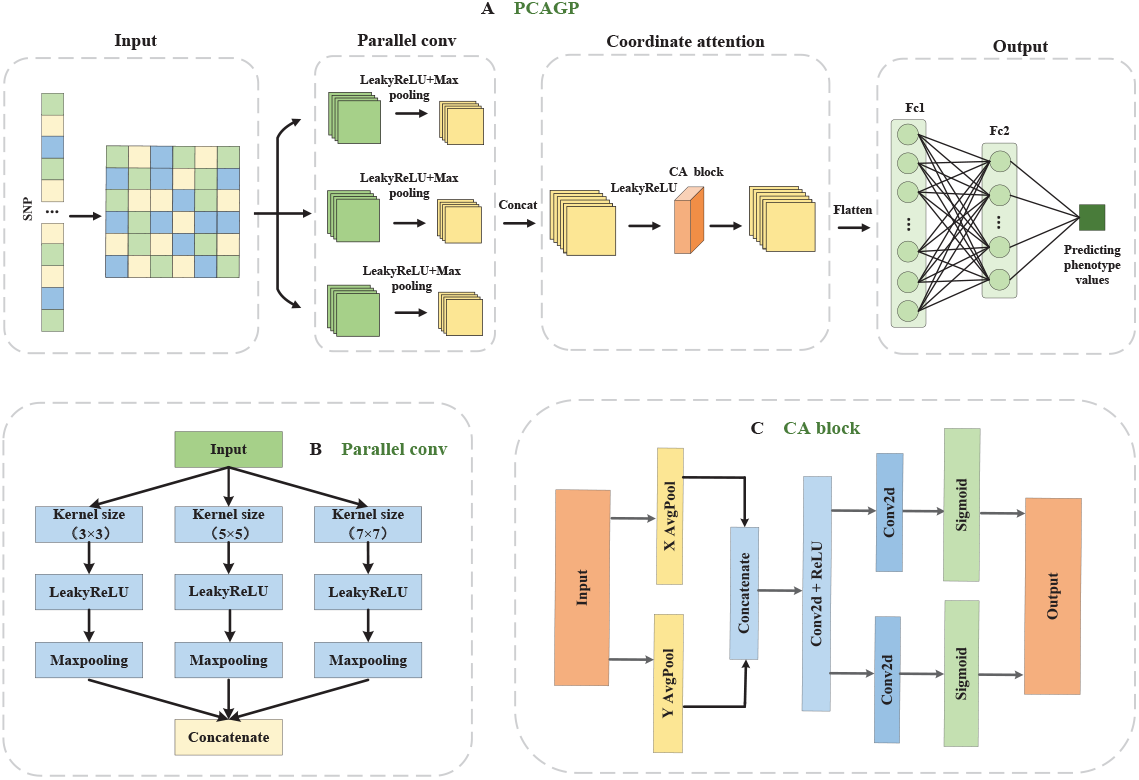
The architecture of PCAGP. (**A**) The flow chart of PCAGP. This model converts the SNP loci corresponding to each sample into an M×M genotype matrix in the input layer. Then, through the parallel convolution layer, three (3×3, 5×5, 7×7) convolution kernels are used for feature extraction. Next, through the coordinate attention layer, the inter-channel and spatial location information are effectively utilized. Finally, through the fully connected layer, the predicted value of the phenotype is output. (**B**) Parallel convolution module. (**C**) Block diagram of the CA attention module. Firstly, the input data is averaged pooled in horizontal and vertical directions. Then, the pooled data is concatenated in the channel dimension. Next, the 1×1 convolution is used to compress the number of channels, and the complete feature vector is re-divided into horizontal and vertical direction vectors, and then the number of channels of the feature vectors in the two directions is readjust by 1×1 convolution. Finally, the Sigmoid function was used to weight the original input data in two directions.

The input of the PCAGP model is the label encoding of the genotype data after screening. And the encoded SNP loci corresponding to each sample are converted into a matrix of size *M* × *M* as the network input.

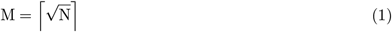

In equation (1), the symbols ⌈ ⌉ represent the ceiling function, where *N* denotes the number of SNP loci corresponding to each sample.

This model performs feature extraction through a parallel convolution module, which can extract information from original data more effectively, as shown in Figure 1B. Moreover, a coordinate attention mechanism is added after convolution to assign different weights to the features, as shown in Figure 1C.

The input first passes through three parallel convolution layers, each of which uses convolution kernels of different sizes, namely 3×3, 5×5, and 7×7, to capture the different-scale features of the input data. Among them, the first convolution layer uses 3×3 convolution kernels with a stride size of 1 and a fill of 1. The second convolution layer uses 5×5 convolution kernels with a stride size of 1 and a fill of 2. The third convolution layer uses 7×7 convolution kernels with a stride size of 1 and a fill of 3. The feature mapping of these three convolution modules is expanded from 1 channel to 16 channels. Fillers of different sizes ensure consistent output dimensions, and a 4×4 max pooling layer is used to reduce the size of the feature map. Then, feature fusion is performed and the output of the parallel convolution module is concatenated in the channel dimension to form a new feature matrix. Subsequently, activation is carried out through the dropout layer (dropout=0.2) and the Leaky ReLU activation function to prevent overfitting from increasing the non-linear representation ability. Subsequently, the coordinate attention module is connected and the spliced feature map is input into the CA block to generate directional attention weights. The feature map is weighted to enhance the expression of effective features. Finally, the feature map processed by the coordinate attention mechanism is flattened into a 1D vector, and the predicted value of the phenotype is output through two fully connected layers.

The CA block is a module based on coordinate attention, designed to enhance the model’s spatial feature representation capability (Hou et al., 2021). Its design integrates attention mechanisms in both the horizontal and vertical directions, while also considering the information between feature channels. The module first extracts global information in the horizontal and vertical directions through two adaptive average pooling operations. Then, it concatenates these two feature maps along the channel dimension and reduces the dimensionality of the concatenated features using a 1×1 convolution layer, batch normalization (BN) layer, and the ReLU activation function. Afterwards, the dimension-reduced features are fed into two separate 1×1 convolution layers, which generate horizontal and vertical attention weights through the Sigmoid function. Finally, the horizontal and vertical attention weights are multiplied by the original feature maps to obtain the weighted output features.

In the PCAGP model, a batch normalization layer is also included (Bjorck et al., 2018). By normalizing the data to a zero-mean and unit-variance distribution, this method helps prevent overfitting and gradient instability, while also improving training efficiency in parameter optimization. For a given feature *x*, the batch normalization formula is as follows:

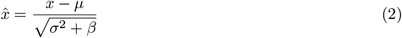

In equation (2), *µ* represents the mean of the features, *σ*^2^ denotes variance of the features, and *β* represents a small constant added to prevent division by zero.

We use the Leaky ReLU activation function (Xie et al., 2020), which is an improved version of the ReLU activation function. Adjusts the negative region of the traditional ReLU activation function, can better maintain the flow of information in negative areas, and effectively alleviates the problem of vanishing gradients. The expression for the Leaky ReLU activation function is as follows:

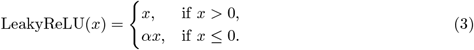

In equation (3), *α* is a small constant, typically set to 0.01.

Due to the challenge of the problem of a large number of SNPs and a small sample size in GS (Wang et al., 2023). This can lead to an overfitting issue that affects the stability and predictive performance of the model. Regularization is a common solution in machine learning to prevent overfitting. Its core function is to limit the degree of freedom of the model parameters by adding additional penalty terms. It improves the generalization ability, enhances the stability of the model, and realizes feature selection to some extent. In the PCAGP model, to mitigate the impact of outliers on training results, we use a loss function that includes the Huber loss function (Meyer, 2021) and an L2 regularization term. The loss function used in the PCAGP model is represented as follows:

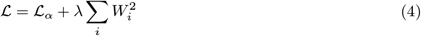

In equation (4), *ℒ*_*α*_ represents the Huber loss function, **W** denotes the weight vector in the model, and *λ* is the regularization coefficient controlling the degree of regularization. This means that larger weights in the model are more penalized, guiding the model to prefer smaller weight values.

The huber loss function is a combination of mean squared error (MSE) and mean absolute error (MAE), and it is more robust than MSE when there are outliers in the data. It applies MSE for small errors and MAE for large errors, providing better performance when handling outliers. The huber loss function is expressed as follows:

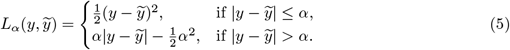

In equation (5), *y* represents the true value, *y* represents the predicted value, and *α* is a hyperparameter used to control the error threshold.

### Experiment setup

We use 5-fold cross-validation to evaluate the prediction performance of the model (Fushiki, 2011). This evaluation method divides the dataset into five equal-sized subsets. In each iteration, four subsets are used for training, while the remaining one is used for validation. This process is repeated five times, and the final performance is evaluated by calculating the average of the evaluation metrics. This method minimizes the risk of overfitting, makes full use of the data, and ensures a more stable performance evaluation. We conducted experiments using the PCAGP model on the four datasets mentioned above and compared performance using two evaluation metrics: the Pearson correlation coefficient (PCC) and mean squared error (MSE). The PCC measures the linear relationship between two variables and has a range of [-1, 1]. The prediction accuracy mentioned in this study refers specifically to PCC values. The MSE quantifies the difference between the predicted values and the true values. MSE values are always nonnegative, with lower values indicating better model prediction performance. All experiments were performed on a computer equipped with an Intel i7 CPU and an NVIDIA GeForce RTX 4090 GPU, using a model learning rate of 0.00009 and running for 150 epochs per iteration.

## Results

### Comparison with other prediction methods

To evaluate the performance of PCAGP in genome prediction compared to traditional method (GBLUP), two machine learning approaches (RF, SVR), and three deep learning models (DeepGS, DNNGP, and SoyDNGP), we performed precision analyzes for different agronomic traits on four datasets of wheat, soybean, and rice. The settings of these models are as follows: for RF, we set the maximum depth of the trees (max depth) to 5, and the number of trees in the forest (n estimators) to 50. For DeepGS and DNNGP, we recreated their model architectures according to the details provided in the original research literature using Python. For the SoyDNGP model, we directly cited the results of the soybean13784 dataset in the original literature. The remaining models used their default parameters as defined in their respective libraries.

The results of the prediction accuracy between PCAGP and other methods on the soybean13784 dataset are shown in Figure 2. The prediction accuracy of the four traits is higher than that of the other five methods. Among all prediction methods, the agronomic trait of SdWgt shows the highest prediction accuracy. Although for the oil trait, PCAGP (0.784) performs slightly worse than GBLUP (0.787), the prediction accuracy for other traits such as poretin, SdWgt, and Yield improved by 0.4%, 0.1%, and 1%, respectively. Based on the average prediction accuracy for all traits, RF was the worst method (0.7) and PCAGP the best (0.796). GBLUP was the second-best method (0.794). PCAGP outperformed GBLUP, RF, SVR, DeepGS, DNNGP, and SoyDNGP by 0.4%, 13.7%, 2.7%, 5.1%, 1.8%, and 5.4%, respectively. We also compared the MSE values of different methods in Table 1. On average, the MSE value of the PCAGP model was 30.9%, 8.5%, 8.9%, 92.2%, and 20.8% lower than RF, SVR, DeepGS, DNNGP, and SoyDNGP, respectively. This indicates that DNNGP still has deficiencies in the accuracy prediction of quantification.

**Table 1:** MSE of seven methods on the soybean13784 dataset.

| Trait | GBLUP | RF | SVR | SoyDNGP | DNNGP | DeepGS | PCAGP |
| --- | --- | --- | --- | --- | --- | --- | --- |
| Protein | **0.0052** | 0.0066 | 0.0057 | 0.0061 | 0.0963 | 0.0063 | 0.0053 |
| Oil | **0.0049** | 0.0066 | 0.0055 | 0.0059 | 0.1163 | 0.0059 | 0.0051 |
| SdWgt | **0.0034** | 0.0077 | 0.0047 | 0.0057 | 0.0154 | 0.0046 | 0.0035 |
| Yield | 0.0111 | 0.0147 | 0.0110 | 0.0134 | 0.0874 | 0.0102 | **0.0107** |

Note: Bold values indicate the best performance for each trait.

**Figure 2.**
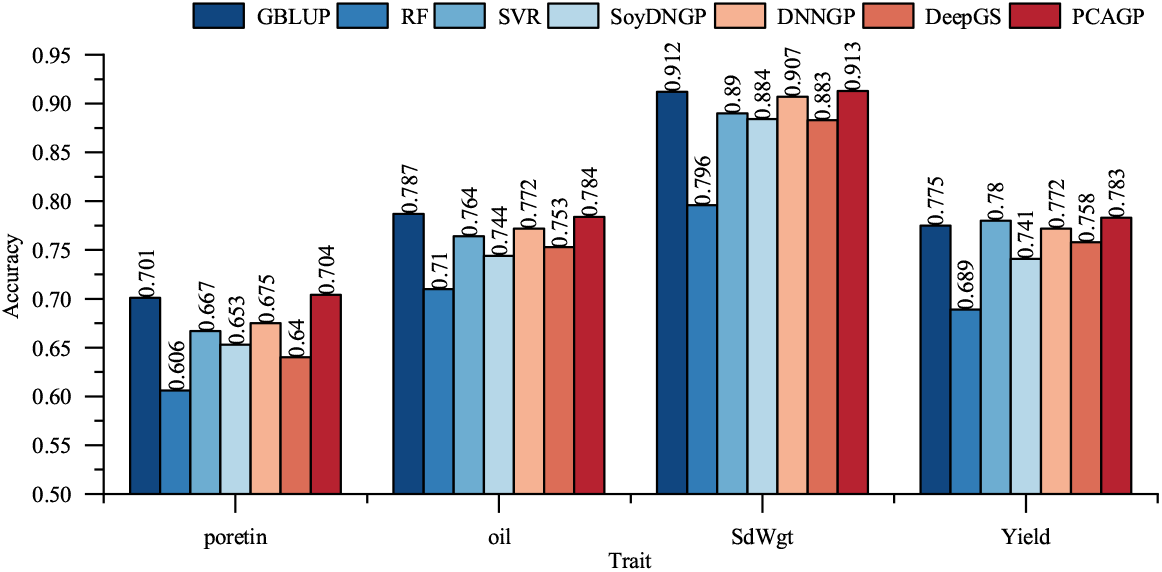
Prediction accuracy of seven methods on the soybean13784 dataset. Protein, oil, SdWgt, and Yield represent protein content, oil content, hundred-seed weight, and yield. GBLUP, genomic best linear unbiased prediction; RF, random forest; SVR, support vector regression; SoyDNGP, a network deep learning framework genomic prediction in soybean breeding; DNNGP, deep neural network for genomic prediction; DeepGS, genomic prediction model based on deep learning framework; PCAGP, parallel convolutional attention network for crop genomic prediction.

As shown in Figure 3, the prediction accuracy for four agronomic traits on the soybean573 dataset is higher than that of the other six methods. We improved by 4.3%, 2%, 2.4%, and 6.1%, respectively, with the second-best forecasting method (GBLUP). The comparison reveals that DeepGS performs the worst, particularly in predicting the average 100-grain weight for the years Yield 13, Yield 14, Yield 17, and Yield 18, where it fails to make predictions. This indicates that the model has poor generalization ability and an unstable training process. Based on the average prediction accuracy for all traits, PCAGP performed the best (0.809), and PCAGP outperformed GBLUP, RF, SVR, DNNGP, and SoyDNGP by 3.7%, 5.1%, 8.5%, 8.4%, and 33.1%, respectively. Furthermore, for the difference between the predicted and true values of PCAGP, the accuracy of the trait prediction of our method was higher than that of the other six methods. The MSE values of different methods are shown in Table 2. On average, the MSE value of the PCAGP model decreased by 10.9%, 18.2%, 25%, 62.2%, 47.1%, and 50%, respectively, compared to GBLUP, RF, SVR, DeepGS, DNNGP, and SoyDNGP.

**Table 2:**
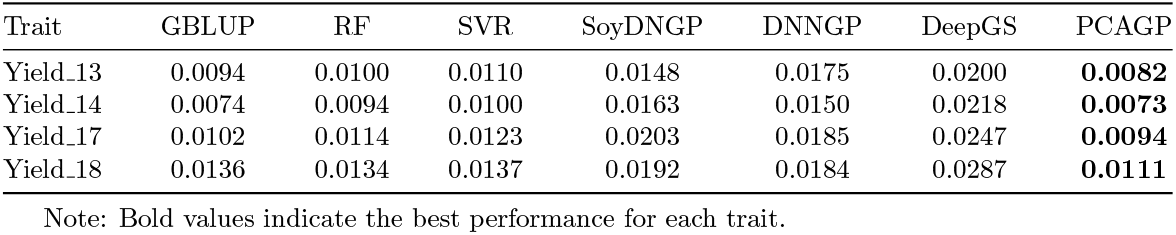
MSE of seven methods on the soybean573 dataset.

**Figure 3.**
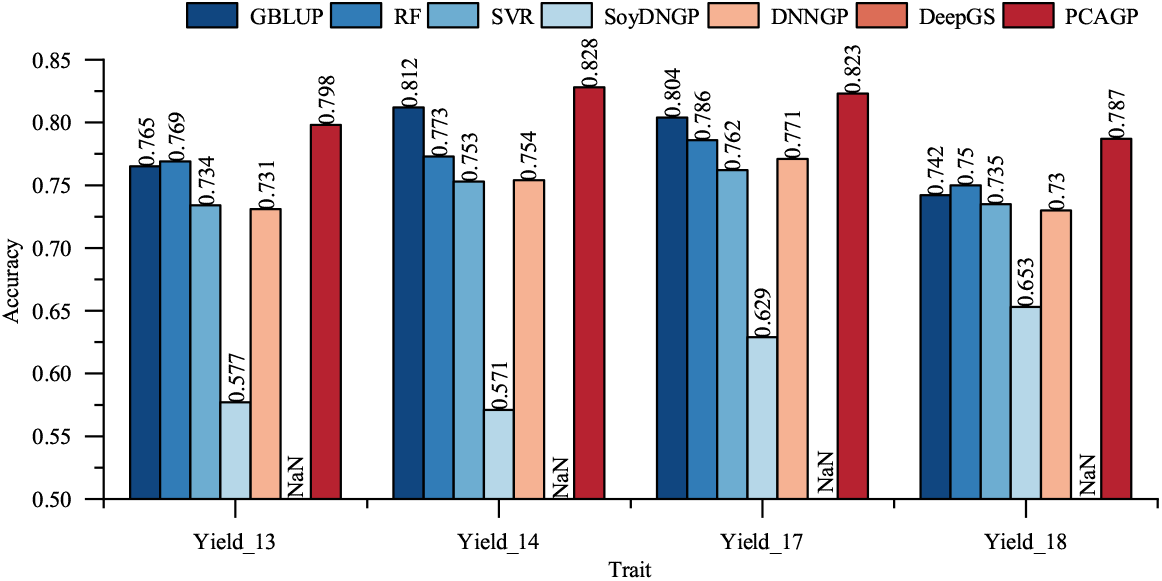
Prediction accuracy of seven methods on the soybean573 dataset. Yield 13, Yield 14, Yield 17, and Yield 18 represent average hundred-seed weight in 2013, average hundred-seed weight in 2014, average hundred-seed weight in 2017 and average hundred-seed weight in 2018.

The experimental results on the wheat2000 dataset are shown in Figure 4. The prediction accuracy of the six traits is higher than that of the other six methods. Among all prediction methods, the agronomic trait of GL (0.74) shows the highest prediction accuracy. We improved by 0.6%, 2.8%, 2.8%, 1.9%, 3.2%, and 2.9%, respectively, with the second-best forecasting method (DNNGP). On average, PCAGP outperforms the other six methods to varying degrees in predicting all traits. Based on the average prediction accuracy for all traits, PCAGP outperformed GBLUP, RF, SVR, DeepGS, DNNGP, and SoyDNGP by 5%, 7.9%, 5%, 26.7%, 2.4%, and 14.1%, respectively. We compared the MSE values of different methods in Table 3. On average, the MSE value of the PCAGP model was 6.2%, 11.6%, 8.7%, 12.7%, 87.7%, and 18.5% lower than GBLUP, RF, SVR, DeepGS, DNNGP, and SoyDNGP, respectively.

**Table 3:** MSE of seven methods on the wheat2000 dataset.

| Trait | GBLUP | RF | SVR | SoyDNGP | DNNGP | DeepGS | PCAGP |
| --- | --- | --- | --- | --- | --- | --- | --- |
| TKW | 0.0260 | 0.0262 | 0.0248 | 0.0298 | 0.0958 | 0.0264 | **0.0235** |
| TW | 0.0134 | 0.0137 | 0.0138 | 0.0149 | 0.1727 | 0.0148 | **0.0124** |
| GW | 0.0099 | 0.0106 | 0.0106 | 0.0118 | 0.1339 | 0.0110 | **0.0096** |
| GL | **0.0053** | 0.0065 | 0.0071 | 0.0066 | 0.0686 | 0.0062 | 0.0055 |
| GH | 0.0206 | 0.0238 | 0.0216 | 0.0243 | 0.1022 | 0.0223 | **0.0193** |
| GP | 0.0125 | 0.0122 | 0.0123 | 0.0138 | 0.0927 | 0.0134 | **0.0117** |

Note: Bold values indicate the best performance for each trait.

**Figure 4.**
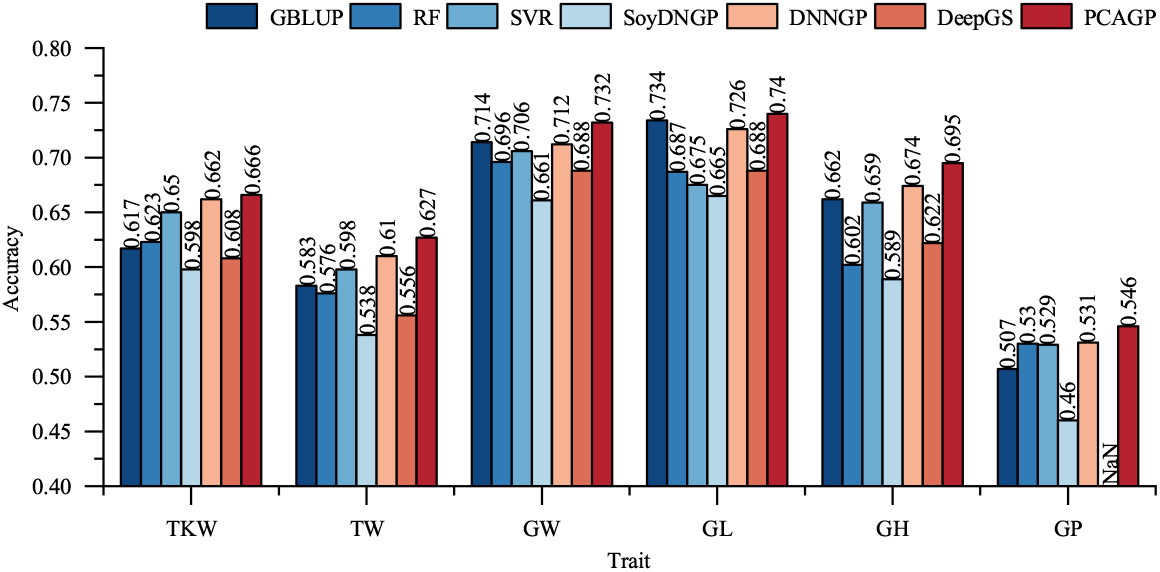
Prediction accuracy of seven methods on the wheat2000 dataset. TKW, TW, GW, GL, GH, and GP represent thousand kernel weight, test weight, grain width, grain length, grain hardness, and grain protein.

As shown in Figure 5, on the rice529 dataset, we compared the prediction accuracy of the other six methods for four agronomic traits: stem dry weight (SDW), stem fresh weight (SFW), grain length (GL) and the ratio of the total projection area to the bounding box area (TBR R). PCAGP outperformed other methods, and compared to SVR, it improved by 2.5%, 3.4%, 3.8%, and 4.4%, respectively. Based on the average prediction accuracy for all traits, GBLUP was the worst (0.426) and PCAGP the best (0.651). PCAGP outperformed GBLUP, RF, SVR, DNNGP, and SoyDNGP by 53.1%, 24.8%, 3.5%, 8.3%, and 21%, respectively. The MSE values of different methods are shown in Table 4. On average, the MSE value of the PCAGP model was 32.6%, 26.2%, 13.9%, 26.2%, 94.5%, and 22.5% lower than GBLUP, RF, SVR, DeepGS, DNNGP, and SoyDNGP, respectively.

**Table 4:** MSE of seven methods on the rice529 dataset.

| Trait | GBLUP | RF | SVR | SoyDNGP | DNNGP | DeepGS | PCAGP |
| --- | --- | --- | --- | --- | --- | --- | --- |
| SDW | 0.0294 | 0.0218 | 0.0170 | 0.0205 | 0.2800 | 0.0251 | **0.0158** |
| SFW | 0.0193 | 0.0165 | 0.0131 | 0.0159 | 0.3944 | 0.0215 | **0.0119** |
| GL | 0.0179 | 0.0200 | 0.0189 | 0.0181 | 0.1611 | 0.0178 | **0.0152** |
| TBR.R | 0.0267 | 0.0250 | 0.0213 | 0.0237 | 0.2951 | 0.0207 | **0.0191** |

Note: Bold values indicate the best performance for each trait.

**Figure 5.**
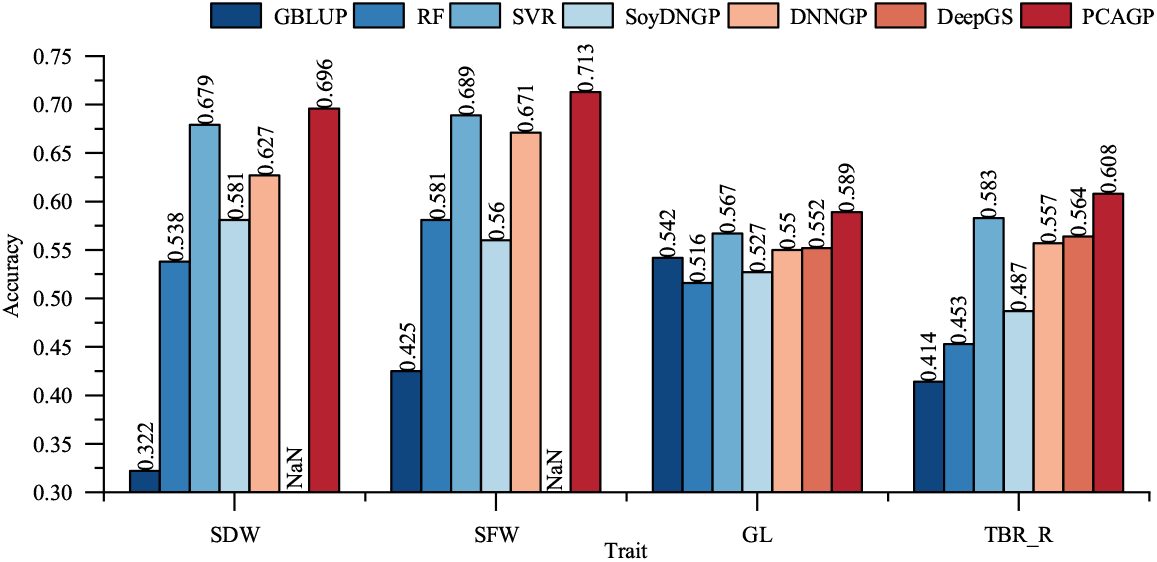
Prediction accuracy of seven methods on the rice529 dataset. SDW, SFW, GL, and TBR R represent stem dry weight, stem fresh weight, grain length, and the ratio of total projected area to bounding rectangle area.

The four datasets used in this study exhibit a significant variation in sample size, with the largest sample size consisting of 13784 individuals and the smallest having only 529 individuals. Although the sample size was relatively small, PCAGP still maintained high precision of genomic prediction, which demonstrated the advantages of the PCAGP model as a genomic prediction tool.

In summary, compared to popular models such as GBLUP, RF, SVR, DeepGS, DNNGP, and SoyDNGP, the PCAGP model demonstrates greater prediction accuracy. Moreover, the PCAGP model is not limited to predicting the phenotype of one crop, but can accurately predict the phenotype values of different crops and has good applicability.

### Effects of sample size on prediction methods

The selection of the size of the training population represents one of the most critical factors that affect the precision of genomic prediction (Xavier et al., 2016), as it directly influences predictive performance. A reasonable size training population can capture the genetic variation of the target trait to the greatest extent, thus improving the reliability and stability of the prediction. To systematically assess the effects of sample size on the accuracy of phenotypic prediction, we performed comparative analyzes using six traits from the wheat2000 dataset (Crossa et al., 2016). Our experimental design used random sampling to generate subsets of 100, 500, 1000, 1500, and 2000 samples. For each subset size, we implemented five-fold cross-validation, with final performance metrics calculated as averages across all validation folds.

The MSE results of each trait corresponding to different sample sizes are shown in Table 5. With an increase in the sample size, the discrepancy between the predicted and actual values gradually decreases. Figure 6 shows the prediction accuracy of PCAGP with different sample sizes on the wheat2000 dataset. When the sample size increased from 100 to 2000, the prediction accuracy of six traits TKW, TW, GW, GL, GH, and GP increased by 28.1%, 22.7%, 34.3%, 42.9%, 36.3%, and 31.3%, respectively. Although the prediction accuracy of different traits decreases with the decrease in sample size, even with a small sample size of only 100, the average prediction accuracy of our model still exceeded 0.5. The results show that the PCAGP model can obtain good prediction performance even when the sample size is small.

**Table 5:** MSE of PCAGP with different sample sizes on the wheat2000 dataset.

| Trait | PCAGP(100) | PCAGP(500) | PCAGP(1000) | PCAGP(1500) | PCAGP(2000) |
| --- | --- | --- | --- | --- | --- |
| TKW | 0.0411 | 0.0321 | 0.0286 | 0.0278 | 0.0243 |
| TW | 0.0174 | 0.0149 | 0.0148 | 0.0126 | 0.0130 |
| GW | 0.0167 | 0.0139 | 0.0137 | 0.0112 | 0.0098 |
| GL | 0.0097 | 0.0094 | 0.0081 | 0.0063 | 0.0069 |
| GH | 0.0311 | 0.0267 | 0.0235 | 0.0231 | 0.0196 |
| GP | 0.0177 | 0.0172 | 0.0134 | 0.0150 | 0.0117 |

**Table 6:** MSE of different methods under various P value thresholds on the wheat2000 dataset.

| Trait | P value threshold | GBLUP | RF | DNNGP | PCAGP |
| --- | --- | --- | --- | --- | --- |
| TKW | all | 0.0261 | 0.0262 | 0.0958 | 0.0235 |
|  | 0.05 | 0.0241 | 0.0265 | 0.0975 | 0.0228 |
|  | 0.01 | 0.0241 | 0.0265 | 0.0881 | 0.0222 |
|  | 0.001 | 0.0240 | 0.0264 | 0.0753 | 0.0233 |
|  | 0.0001 | 0.0238 | 0.0263 | 0.0554 | 0.0228 |
| TW | all | 0.0134 | 0.0137 | 0.1727 | 0.0124 |
|  | 0.05 | 0.0120 | 0.0138 | 0.1770 | 0.0121 |
|  | 0.01 | 0.0122 | 0.0138 | 0.1685 | 0.0124 |
|  | 0.001 | 0.0123 | 0.0138 | 0.1632 | 0.0129 |
|  | 0.0001 | 0.0126 | 0.0138 | 0.1519 | 0.0127 |
| GW | all | 0.0106 | 0.0106 | 0.1339 | 0.0096 |
|  | 0.05 | 0.0095 | 0.0107 | 0.1343 | 0.0096 |
|  | 0.01 | 0.0096 | 0.0107 | 0.1261 | 0.0095 |
|  | 0.001 | 0.0092 | 0.0107 | 0.1215 | 0.0094 |
|  | 0.0001 | 0.0090 | 0.0113 | 0.1102 | 0.0091 |
| GL | all | 0.0053 | 0.0065 | 0.0686 | 0.0055 |
|  | 0.05 | 0.0050 | 0.0065 | 0.0679 | 0.0053 |
|  | 0.01 | 0.0052 | 0.0065 | 0.0605 | 0.0052 |
|  | 0.001 | 0.0052 | 0.0065 | 0.0527 | 0.0055 |
|  | 0.0001 | 0.0050 | 0.0063 | 0.0486 | 0.0053 |
| GH | all | 0.0206 | 0.0238 | 0.1022 | 0.0193 |
|  | 0.05 | 0.0197 | 0.0238 | 0.0972 | 0.0188 |
|  | 0.01 | 0.0201 | 0.0239 | 0.0922 | 0.0194 |
|  | 0.001 | 0.0201 | 0.0239 | 0.0902 | 0.0195 |
|  | 0.0001 | 0.0198 | 0.0238 | 0.0886 | 0.0197 |
| GP | all | 0.0125 | 0.0122 | 0.0927 | 0.0117 |
|  | 0.05 | 0.0107 | 0.0120 | 0.0867 | 0.0114 |
|  | 0.01 | 0.0112 | 0.0120 | 0.0701 | 0.0113 |
|  | 0.001 | 0.0111 | 0.0121 | 0.0456 | 0.0111 |
|  | 0.0001 | 0.0112 | 0.0121 | 0.0263 | 0.0114 |

**Figure 6.**
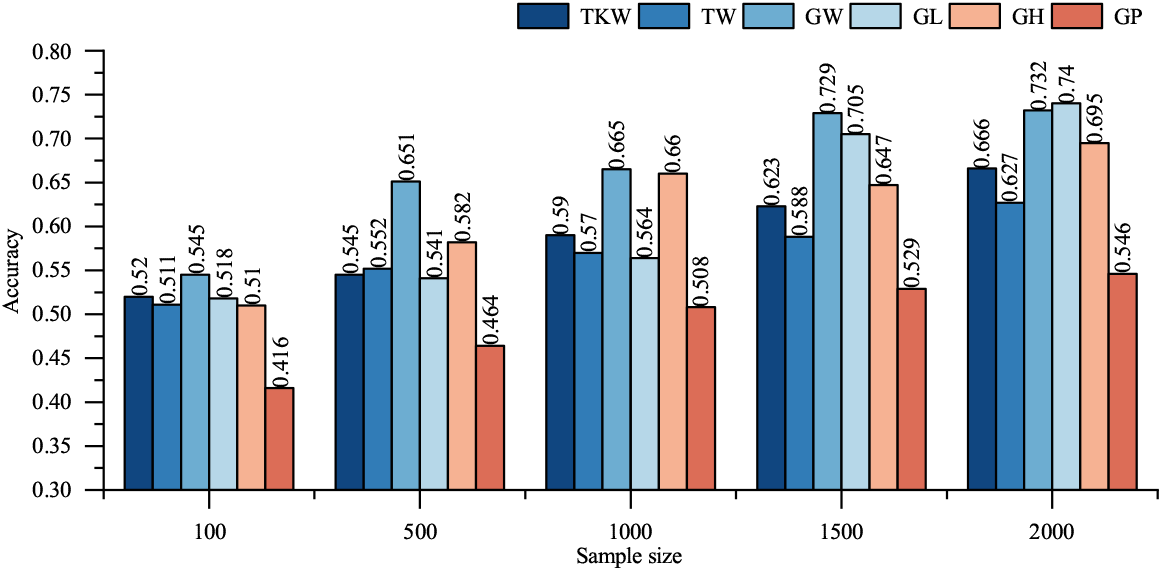
Prediction accuracy of PCAGP with different sample sizes on the wheat2000 dataset. TKW, TW, GW, GL, GH, and GP represent thousand kernel weight, test weight, grain width, grain length, grain hardness, and grain protein.

### Effects of SNP number on prediction methods

A smaller number of SNPs that are significantly associated with target traits can achieve predictive performance similar to that of unfiltered genome-wide SNPs (Wang et al., 2023). This is because these selected SNPs could capture more accurately the genetic variation related to the trait, thereby improving the prediction accuracy of the model. To study the effects of SNP number on the prediction accuracy of different prediction methods, we used the wheat2000 dataset to compare the prediction results of the GBLUP, RF, DNNGP, and PCAGP models with different p value thresholds (p = 0.05, 0.01, 0.001, and 1e-04) for SNPs selection.

As shown in Figure 7, the results of different SNP numbers show the degree of variation in prediction accuracy within the PCAGP model. We can see that the prediction accuracy of all methods did not change significantly as p-value screening became more rigorous. The results demonstrate that the use of trait-associated SNPs achieves predictive performance similar to that obtained by employing unfiltered genome-wide SNPs. Therefore, it is feasible to screen SNPs using reasonable methods, so that the selected SNPs can reliably predict traits.

**Figure 7.**
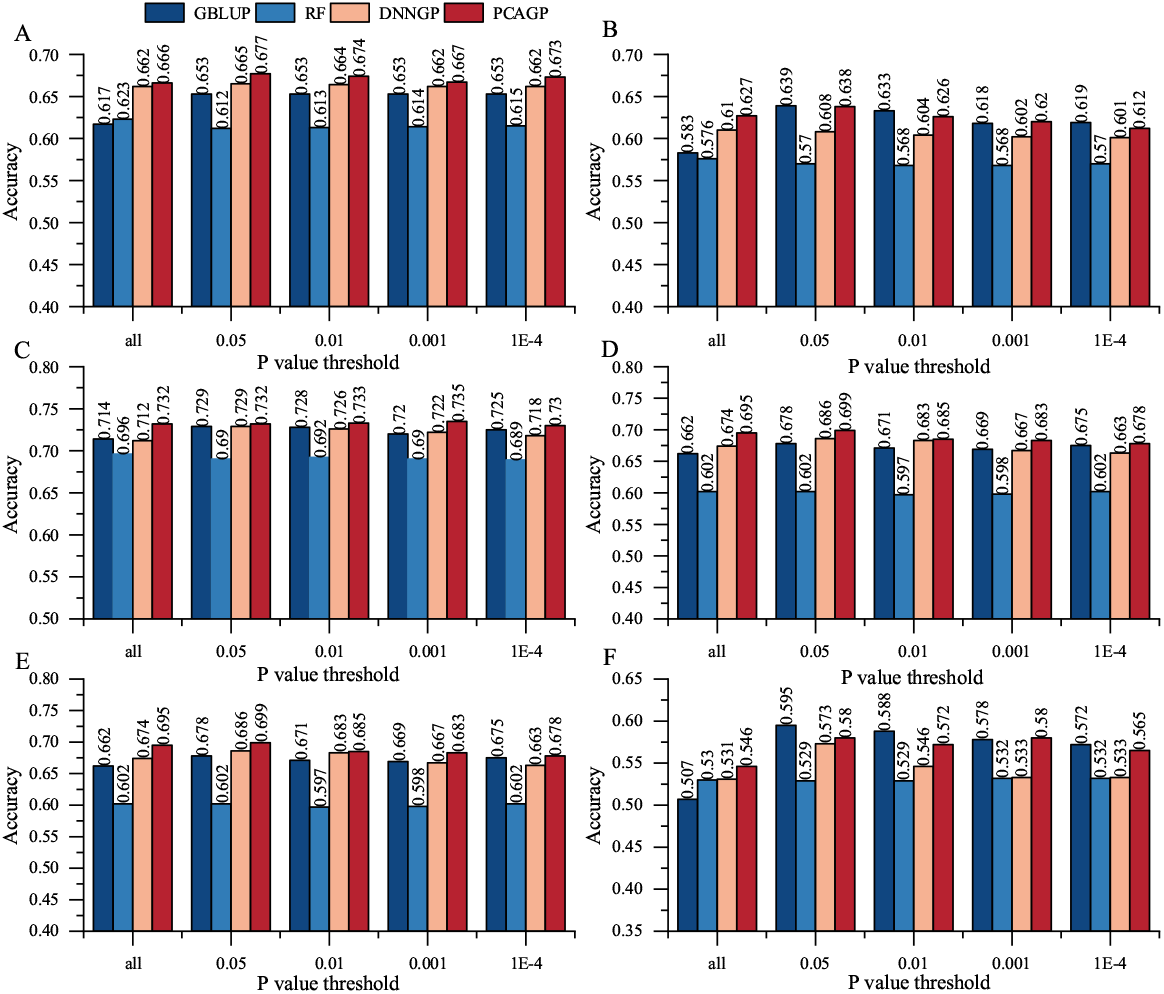
Prediction accuracy of models built using four methods and filtered SNP markers from the wheat2000 dataset. (A) thousand kernel weight (TKW); (B) test weight (TW); (C) grain width (GW); (D) grain length (GL); (E) grain hardness (GH); (F) grain protein (GP).

In addition, reducing the number of input features can reduce computational complexity and effectively prevent overfitting. However, the effectiveness of this approach depends heavily on the accuracy of identifying SNPs that are truly associated with the target traits during the selection process. Improper selection may overlook important genetic signals, which adversely affects the predictive performance of the model. Therefore, a reasonable screening method is needed to select the appropriate number of SNPs, and the selected SNPs could meet the representative characteristics.

## Discussion

The innovation of theoretical methods has laid the foundation for the development of GS technology. However, to truly apply it to breeding, efficient prediction of crop genotypes and phenotypes is essential. There is an urgent need to develop GS methods with greater prediction accuracy. Although models such as GBLUP (VanRaden, 2008), RF (Adetunji et al., 2022), SVR (Maenhout et al., 2007), DeepGS (Ma et al., 2018), DNNGP (Wang et al., 2023), and SoyDNGP (Gao et al., 2023) have made significant contributions to this field, they also exhibit certain limitations. In particular, the classical deep learning models all use a serial structured convolution kernel to extract features. Although it is possible to explore the complex relationships in SNPs to some extent, determining the size of the convolution kernels is also a very time-consuming task. Moreover, the serial structure may also have the risk of information loss. This kind of loss of information can hinder the model’s ability to capture complex genetic interactions and their effects on traits, especially for highly nonlinear genomic data. We propose a method based on a parallel convolutional attention network for crop genomic prediction (PCAGP), which can effectively learn the relationships among genotype data through parallel convolutions to predict phenotypes. We apply it to different crops to predict traits and compare the prediction accuracy of different models, which outperforms current genomic selection models. The main advantages of our model are as follows.

Firstly, the PCAGP model adopts parallel convolution layer simultaneously to extract features, which offers significant computational efficiency advantages. Compared to a serial convolution layer, parallel processing enables significant improvements in computational efficiency through hardware acceleration, allowing faster analysis of large-scale data. This makes the parallel module suitable for processing large-scale genomic data.

Secondly, the PCAGP model introduces a coordinate attention module, which ensures that the genotype data features extracted through parallel convolution are assigned different weights based on the correlation of channel and spatial positional information. This enables the model to exhibit higher prediction accuracy.

Finally, the overall design of the PCAGP model does not rely on deep network layers. In phenotypic prediction, the number of SNPs is much greater than the number of samples, and the use of deep network layer may lead to overfitting. A wider network layer is more beneficial for feature representation, while minimizing information loss and preserving more data details.

## Conclusion

This paper introduces a new genome-wide prediction method called PCAGP. This model transforms the SNP sites of each sample into a two-dimensional matrix representation. Using a parallel convolution architecture, it simultaneously extracts genotype features using convolution kernels of varying sizes to achieve better feature extraction. Incorporating a coordinate attention mechanism enables precise acquisition of localization information while effectively capturing long-range dependencies. Experimental results demonstrate that our model exhibits strong generalization ability across diverse crop species for agronomic trait prediction tasks, consistently achieving high prediction accuracy. These findings validate the robustness and effectiveness of the PCAGP model, suggesting its broad potential for application in GS research and breeding programs.

## Author contributions statement

W.P. performed the experiments and drafted the manuscript. Y.S analyzed the data. Y.S, Y.Z., and L.C. revised the manuscript. All authors read and approved the final manuscript.

## Acknowledgments

We thank Prof. Lizhong Xiong of Huazhong Agricultural University for sharing rice dataset; we are also grateful to Prof. Xutong Wang of Huazhong Agricultural University for providing soybean dataset.

## Data availability

The implementation code for PCAGP models along with example datasets is accessible at https://github.com/Crop-breeding/PCAGP.

